# Awareness is necessary for predictive learning and prediction-based motor attenuation

**DOI:** 10.64898/2025.12.01.691731

**Authors:** Lexin Liang, Christy K. W. Tam, Evan J. Livesey, Dominic M. D. Tran

**Affiliations:** The University of Sydney, Camperdown, NSW, Australia

**Keywords:** Predictive learning, Prediction attenuation, Prediction error, Expectation violation, Awareness, Motor system, Transcranial magnetic stimulation

## Abstract

Predictive coding theories posit that the brain continuously updates internal models of the world to anticipate sensory input. Attenuation of neural responses to predictable sensory stimuli is thought to aid in this process of prediction error detection and model updating. The topic of whether predictive processes such as sensory attenuation can occur in minimally conscious states has been hotly debated. The current study provides a novel test for the role of awareness in prediction-based attenuation using a transcranial magnetic stimulation (TMS) prediction task, whereby predictable stimulation is known to lead to attenuated motor system excitability. The experiment tested whether motor attenuation can occur without awareness using a dual task paradigm to mask a predictive cueing relationship. Participants simultaneously completed an n-back task and a TMS-counting task; critically, a visual cue (in the n-back task) consistently predicted the occurrence of a TMS pulse (in the counting task). Instruction conditions manipulated how much participants were informed of this relationship, and awareness was assessed in a post-experimental questionnaire. The results reveal that motor attenuation was present only in participants instructed about the relationship, or in those who became aware of the cue-TMS association. Implicit, predictable cue-TMS exposure was not sufficient to elicit motor attenuation in unaware participants. These findings show that awareness of the relationship is necessary for predictive learning and prediction-based attenuation in the motor system, challenging the assumption that predictive coding operates automatically.

## Sensory attenuation

It has been widely documented that prediction influences the way our brain perceives and interprets information. For example, when we can predict the occurrence of sensory signals, our neural response is often suppressed compared to signals that are unpredictable. We refer to this down-weighting of sensory information as prediction-based sensory attenuation (e.g. Ford et al., 2007). Sensory attenuation has been proposed to confer an evolutionary advantage since there is a greater need to pay attention to unexpected signals that may indicate a threat, whilst less attention can be diverted to predictable signals to conserve energy (Crapse & Sommer, 2008). The corollary discharge theory of sensory attenuation explains how the brain distinguishes between sensory inputs caused by external stimuli and those resulting from self-generated actions (Blackmore, Frith & Wolpert, 1999). When the brain sends a motor command to move part of the body (e.g., hand, eyes, limbs, etc.), it simultaneously sends a copy of this command (a corollary discharge) to sensory areas of the brain. If the sensory input matches the predicted outcome of the motor command, the brain attenuates or suppresses the sensory response. Dampening the salience of self-generated sensations is thought to explain why we cannot tickle ourselves (Schafer & Marcus, 1973).

Sensory attenuation can be observed by comparing neurophysiological signals for self-generated sounds and externally generated sounds (e.g. Bäss et al., 2008). Using electroencephalography (EEG), studies have measured event-related potentials (ERPs) in response to auditory tones. These auditory evoked potentials include an N1 component which varies in amplitude based on the loudness of the sound (Mulert et al., 2005; Simmons et al., 2011). Attenuation of the N1 component has been observed when the tone is made predictable, either through self-generation (Elijah et al., 2016) or a visual warning cue (Ford et al., 2007). In a further a study by Elijah et al. (2016), participants generated an auditory tone via a button press and either received the playback immediately or after a delay. Immediate tones produced greater N1 attenuation compared with delayed tones, which can be explained as a bias towards expecting immediate sensory feedback. However, when participants were repeatedly exposed to delayed tones during an initial training phase, the difference in attenuation between immediate and delayed tones was eliminated. This finding demonstrates that it is the familiarity and expectation of the action-feedback relationship, which can be learned, that is critical for the attenuation effect, rather than any differences in temporal characteristics per se.

## Motor attenuation

Recent research has investigated prediction-based attenuation in non-sensory areas (Tran & Livesey, 2021). A study by Tran et al. (2021) used transcranial magnetic stimulation (TMS) to deliver pulses to the primary motor cortex. Stimulation of the motor cortex can elicit a motor-evoked potential (MEP) in the contralateral hand when measured with electromyography. The peak-to-peak amplitude of the MEPs provides an index of the excitability of the motor system. The authors found that predictable TMS pulses (i.e., those that were self-initiated or signaled by a warning cue) produced smaller MEPs compared to unpredictable or unexpected TMS pulses of the same intensity. These prediction-based attenuation effects were also observed when controlling for motor responses or without performing any response. In one experiment, participants passively observed a visual representation of an analogue stopwatch that signalled when to expect TMS stimulation. Greater attenuation was observed to TMS pulses delivered at a predictable timepoint (i.e., on time) compared with those delivered unexpectedly early. While the function of attenuation processes in M1 are yet to be determined (see Tran et al., 2021 for different hypotheses), the results from these experiments show that the predictability of TMS pulses can modulate corticospinal excitability in the motor system and that prediction-based attenuation is not confined to sensory systems.

Together, the evidence of prediction-based attenuation in both the sensory and motor domains suggest that predictive coding may be regulated by domain-general mechanisms. To examine this hypothesis, Tran et al. (2025) used a combined TMS-EEG setup to measure motor attenuation using TMS and simultaneously measure sensory attenuation in response to the sound accompanying TMS when a pulse is delivered. The TMS coil produces a salient auditory coil “click” during stimulation. The magnitude of the sensory and motor attenuation effects varied with the predictability of the stimulation/sound as expected. Moreover, the strength of sensory attenuation to the “click” of the TMS predicted the strength of motor attenuation from direct stimulation of the motor cortex. This close correspondence between prediction-based neural attenuation across the sensory and motor systems suggests that they may be governed by domain-general mechanisms. Tran et al. (2025) observed this relationship despite also finding evidence that it is the prediction of M1 stimulation—and not prediction of salient sensory signals—that drives the MEP attenuation effect.

## Awareness in predictive coding

Predictive coding is thought to be governed by automatic processes updating the generative model of our environment, based on incoming sensory information. When a prediction about sensory input mismatches the external environment, a prediction error is propagated up the hierarchy to prompt higher-level revisions of expectations (Parr et al., 2022). This recursive process of generating prediction errors and updating predictive models is thought to occur in the absence of conscious awareness. Support for this automaticity comes from studies showing neurophysiological evidence of sensory prediction errors in participants with reduced or altered states of consciousness (see Tivadar et al., 2021). One way of studying auditory prediction and prediction error is using an oddball paradigm where a series of identical sounds are presented and are occasionally interrupted with deviant sounds (the *oddball*; Garrido et al., 2009). Neural signatures of auditory prediction can be measured using EEG, such as the mismatch negativity (MMN) signal—a negative EEG component derived from the difference in activity between the standard stimuli and deviant stimuli (Garrido et al., 2009); or the P300—a positive EEG component in response to novel stimuli (Picton, 1992). Repeated auditory stimuli have been shown to produce attenuated neural responses compared with the oddball, which is thought to reflect differences in predictive processes between the two types of stimuli (Summerfield et al., 2008). A meta-analysis by Tivadar et al. (2021) found that EEG signatures such as MMN and P300 were primarily preserved in patients in reduced states of consciousness such as during sleep, anesthesia, and coma, although these signatures were attenuated with decreasing levels of consciousness. These findings were taken as evidence to suggest that predictive learning processes are retained even when consciousness is reduced or absent, suggesting that prediction and prediction error processes occur automatically.

However, the literature on awareness and neurophysiological markers of expectation violation has not always been consistent, particularly when controlling for confounding variables such as adaptation (e.g., Bekinschtein et al., 2009; Nourski et al., 2018). Investigations into the role of awareness in predictive coding have primarily used auditory stimuli as it is thought that sounds can still reach the brain without requiring attention. Since TMS can directly activate corticomotor neurons, the motor attenuation paradigm offers a novel way of testing for the role of conscious awareness in predictive coding. Hence, the current experiment sought to test whether the motor attenuation effects observed by Tran and colleagues (2021; 2025) could be observed without awareness.

Studying awareness in healthy individuals without disorders of consciousness can be tricky. The experiment requires some way of masking a simple predictive relationship between the preceding warning signal (or cue) and the following outcome. Research on conditioning offers valuable insights into how awareness (e.g., Lovibond & Shanks 2002) and automaticity (e.g., Perruchet 2015; Tran et al., 2020) can be tested in predictive learning. A study by Wiedemann et al. (2016) experimentally manipulated awareness in an eyeblink conditioning paradigm to test whether knowledge of the predictive relationship is necessary for conditioning. In the experiment, participants simultaneously completed two tasks: an n-back task where they responded whenever the current shape was the same as the one on the previous trial, and a go/no-go task where they were instructed to respond to a tone and withhold their response to an air puff directed at the participant’s eye. When presented with an air puff, participants will typically produce an unconditioned blink response. However, if participants learn that the air puff is consistently preceded by a predictive cue or warning signal (e.g., a specific shape), participants will start to blink to the predictive cue in advance of the air puff delivery (i.e., eyeblink conditioning). The use of the dual task masked a predictive relationship between a specific shape in the n-back task and the occurrence of the air puff in the go/no-go task. Awareness of this shape-air puff relationship was further manipulated by randomly allocating participants into one of three conditions: informed (instructed about the specific stimulus that predicted the air puff), relational (instructed that there was a relationship between the two tasks but not the nature of the relationship), or uninformed (provided no information about the relationship between the two tasks).

The authors observed differences across the instruction groups: eyeblink conditioning emerged early during the dual task for the informed group, emerged over time for the relational group, and was absent for the uninformed group. That is, despite a specific shape consistently preceding an air puff, if participants did not learn or were unaware of this predictive relationship having compartmentalised the two tasks as separate representations, then participants did not blink to the predictive cue (the shape signaling the arrival of the air puff). Participants were further classified as aware or unaware based on knowledge of the predictive (shape-air puff) relationship examined in a post-experimental questionnaire. Participants classified as aware showed evidence of eyeblink conditioning whilst participants classified as unaware failed to show evidence of conditioning. The strength of this dual task is its ability to manipulate attention to the *relationship* between the stimuli (i.e., the warning cue, and outcome) while ensuring attention to the individual stimuli. These findings suggest that contingency awareness of the predictive relationship between a warning signal (e.g., a shape) and an outcome (e.g., an air puff) play a causal role in learning and conditioning.

## The current study

The current study investigated the role of awareness in prediction-based motor attenuation using TMS. Taking inspiration from Wiedemann and colleagues (2016), participants performed two tasks simultaneously: an n-back task, where they were instructed to respond to repeated images; and a counting task, where they were instructed to count the number of TMS pulses intermixed with auditory tones. The dual task masked a predictive relationship between a specific image in the n-back task and the presentation of a TMS pulse in the counting task.

## Method

### Participants

Participants were Psychology students at The University of Sydney who completed SONA experiments for course credit. Based on prior motor attenuation (Tran et al., 2021) and between-group TMS designs (Tran et al., 2019), we aimed to test 24 participants per condition. Seventy-four participants were recruited with 2 participants withdrawing due to discomfort with the TMS intensity required to elicit an MEP. Participants (Age: M = 20.1, SD = 3.1; Gender: 51 female, 19 male, 2 non-binary) were sequentially allocated to informed (*n* = 24), relational (*n* = 24) and uninformed (*n* = 24) conditions, according to the order in which they participated. All participants completed a TMS safety questionnaire based on Rossi et al. (2009) and provided informed consent before commencing the experiment. The study was approved by the Human Research Ethics Committee of The University of Sydney.

### Apparatus and stimuli

The experiment was run on a Windows 7 PC using PsychoPy2 to control stimulus presentation. Stimuli were displayed on a 24-inch ASUS monitor (1920 × 1080 resolution, 60 Hz refresh rate) at a viewing distance of ∼57 cm.

### EMG

Surface electrodes were attached to the right hand for electromyography (EMG) recording. Skin was prepped via exfoliation using a small scouring pad and a 70% v/v isopropyl alcohol wipe. A pair of Ag/AgCl electrodes were placed on the proximal and distal surfaces of the FDI muscle, and a ground electrode was placed over the ulnar styloid process of the wrist. EMG activity was recorded from 100 ms pre-stimulation to 400 ms post-stimulation. This signal was digitally converted (sampling rate: 4 kHz, bandpass filter: 0.5 Hz to 2 kHz, mains filter: 50 Hz, and anti-aliasing) and stored offline for analysis using LabChart software (Version 8, ADInstruments).

### TMS

Transcranial magnetic stimulation (TMS) was administered using a MagStim 200^2^ stimulator and a 70 mm D70^2^ figure-eight coil. A cotton cap was secured on the participant’s head to help locate the hand region of the left motor cortex. The coil was held tangentially to the scalp with the coil oriented 45° from the midline. Starting at 5 cm lateral and 1 cm anterior to Cz, the coil was moved around to determine the motor cortex “hotspot” until maximal MEP could be elicited in the FDI. Once the hotspot was determined and marked, the participant placed their head on a chin and forehead rest and the coil was fixed into position with an adjustable mechanical arm (Manfrotto). Resting motor threshold (rMT) was defined as the lowest stimulation intensity that produced an MEP of at least 50 µV in 5 out of 10 consecutive trials (Rossini et al., 2015). Stimulation intensity during the experiment was set to 120% of rMT.

### Procedure

Participants first completed a practice session in which they performed each task separately.

### Counting task

Participants were instructed that they would receive a series of stimuli, one at a time. Sometimes the stimulus would be a sound, and sometimes the stimulus would be a TMS pulse. Their task was to count the number of times they received TMS and report it at the end. There were 25 trials in total (10 TMS and 15 auditory tones), presented in random order. The display duration was 1800ms, with a random ITI between 4500 and 5500ms. At the end of the sequence of trials, participants were prompted to enter the number of TMS pulses they received.

### N-back task

In the second task, participants were told that they would see a series of fruit images on the screen, one at a time. They were instructed to press the space key with their left hand whenever the image was the same as that on the previous trial (1-back task). If the image was different, they were instructed to make no response. They were asked to respond as quickly and accurately as possible. On each trial, a 300 x 300 pixel fruit image (*Apple*, *Banana*, *Orange* or *Strawberry*), was presented in the center of the screen. Each fruit was displayed for 1600 ms, during which participants could respond, followed by a non-response interval of 200 ms, and a random ITI between 2700 and 3700 ms. Feedback was provided for each response: “Correct Press” was shown in green for correct responses; “Do not respond” was shown in red following an incorrect response; and “Miss” was shown in red following a missed repeat. There were 40 practice trials with each fruit presented 10 times each. The sequence was pseudorandomized such that ∼25% of trials were repetitions (1-back matches).

### Dual task

Following practice, participants completed the primary task where they performed the n-back and counting task simultaneously (see Figure 1). They received the following instructions:

**Figure 1.**
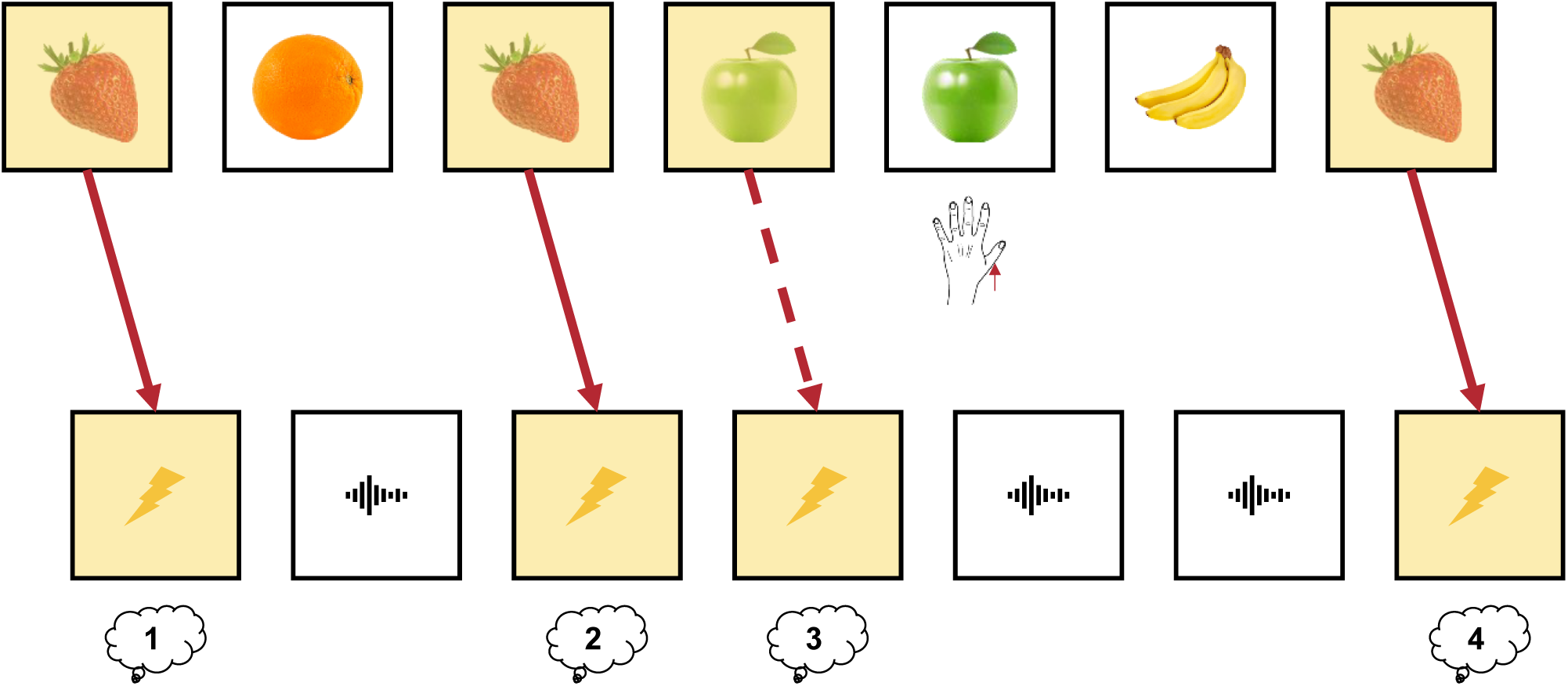
Schematic of the dual task comprising an n-back task with fruit stimuli, and a counting task with TMS single pulses and auditory tones. In the n-back task, participants had to respond when the current stimulus was the same as the one presented previously (i.e., 1-back). In the counting task, participants had to count the number of TMS pulses within a random sequence of TMS and auditory tones. The dual task masked a relationship between the two tasks, whereby one fruit was always followed by a TMS pulse (in the schematic, this is the strawberry; in the experiment, the predictive fruit was counterbalanced across participants). TMS was also sometimes delivered following each of the other fruits. Highlighted squares indicate trials that were followed by TMS: a solid arrow indicates a predictable TMS trial; dashed arrow indicates an unpredictable TMS trial.

*In this experiment, we will assess your ability to switch between two tasks. You need to perform the two tasks in the practice session, but now you will do them simultaneously. You will see a series of images presented on the screen, and you need to press the SPACE key whenever the image is the same as the previous image. In addition, you will also receive a series of sounds and pulses. Please also count the number of TMS pulses you receive and report it at the end of each block*.

Each group then received different instructions to manipulate knowledge of the relationship between the warning cue and TMS pulses: the informed group were told *“Here’s a tip to help you with the combined task: Every time you see the* [one of the fruit images]*, a TMS pulse will be triggered.”*; the relational group were told *“There is a relationship between the two tasks. Working out the relationship will help you in performing the tasks.”*; and the uninformed group were told *“Remember, you need to respond to the repeated fruit and count the number of TMS pulses at the same time.”* There were 5 blocks of the task, with the 1-back and counting task interleaved in a way that each trial presented a fruit image followed by a TMS or tone. Each block presented a sequence of fruit images, with each fruit appearing an equal number of times per block. Trial order was pseudorandomized so that approximately 25% of transitions were one-back repetitions. Immediately following the image presentation, participants received either a TMS pulse or a brief auditory tone. TMS pulses were delivered to coincide with the onset of the post-stimulus interval. One fruit type (predictive cue) was consistently paired with TMS across all its presentations, while the remaining three fruits (non-predictive cue) were paired with TMS on 20% of their trials. The allocation of fruit to the predictive and non-predictive cues was counterbalanced across participants. The number of presentations per fruit within a block varied between 25 and 35 based on the TMS constraints required to establish a predictive relationship with one of the fruits, and a non-predictive relationship with the other three fruits. This resulted in approximately 240–260 TMS trials per participant across five blocks (totaling ∼520–680 trials), with the remaining trials paired with sound. The variability in trial numbers was due to the randomisation of 1-back trials while maintaining the contingency of the predictive (100%) and three non-predictive (25%) cues.

Following the task, participants were asked a series of questions to assess their knowledge of the predictive fruit relationship. The questions increased in specificity about the predictive relationship. First, they were asked an open-ended question *“In the experiment, did you notice any relationship between the fruit images and TMS? If so, what is the relationship?”*. Participants typed their responses in a text box. They were then presented with four rating scales, each accompanied by a fruit image, and were asked to rate how often each fruit was accompanied by TMS on a scale from 1 (*“Never accompanied by TMS”*) to 9 (*“Always accompanied by TMS”*). Finally, they were given a forced choice question *“Which fruit was most often accompanied by TMS?”* and were required to click on one of the four images or type the corresponding number (1-4) to select their answer.

#### Analysis

Motor-evoked potentials for predictable TMS trials were log normalised to the unpredictable TMS trials. MEPs were normalised to reduce large between-participant variability in corticospinal excitability and to normalise a positively skewed distribution. Full reasons for log normalisation are outlined in prior research (e.g., Tran et al., 2021). Log normalised MEPs < 0 indicate smaller predictable MEPs relative to the unpredictable MEPs (i.e., a prediction-based attenuation effect).

First, log normalised MEPs were analysed by instruction condition (Informed, Relational, Uninformed). The Instruction group factor was analysed using a set of orthogonal contrasts comparing: 1) information versus no information [Informed and Relational vs Uninformed: 1 1 - 2]; and 2) full versus partial information [Informed vs Relational: 1 -1 0]. These group contrasts were also analysed in an interaction with block as a linear trend [-2 -1 0 1 2].

The data were also analysed by awareness classification based on the post-experimental questionnaire. Participants were classified aware if they answered *any* of the three questions correctly. That is, they loosely described that TMS always followed one of the fruits (in the free response); rated the predictable fruit as the most likely to be followed by TMS (in the rating judgements); or selected the predictable fruit as the most often to be followed by TMS (in the forced choice). This assessment of contingency awareness meant that participants classified as unaware failed all three questions. While this criterion means that we may classify some participants as “Aware” who were not knowledgeable about the predictive relationship, we can be more confident that any participants classified as “Unaware” have very little knowledge about the predictive relationship.

Second, log normalised MEPs were analysed by awareness classification (Aware, Unware). Awareness classification will also be analysed in an interaction with block as a linear trend [-2 -1 0 1 2].

## Results

Log normalised MEPs for the Informed group were below baseline from block 1 and continued to reduce throughout the blocks (see Figure 2). For the Relational group, log normalised MEPs were around baseline for blocks 1 and 2 and reduced below 0 from blocks 3 to 5. For the Uninformed group, log normalised MEPs remained around baseline for all blocks. Table 1 includes the full list of one-sample t-tests by Group and Block comparing log normalised MEPs against baseline (i.e., the attenuation effect). The contrast comparing information (Informed and Relational) versus no information (Uninformed) showed that overall, across blocks, the information groups had significantly lower log normalised MEPs than the no information group; *t*(69) = 3.89, *p* < .001. This contrast also significantly interacted with Block as a linear trend, indicating that the effect of information on MEPs became larger across blocks; *t*(276) = 2.25, *p* = .025. Comparing full information (Informed) versus partial information (Relational) showed there was an overall benefit of the full instruction, whereby the Informed group had significantly lower log normalised MEPs than the Relational group; *t*(46) = 2.38, *p* = .02. This contrast did not significantly interact with Block as a linear trend, indicating insufficient evidence that the difference between full and partial information on MEPs changed across blocks; *t*(184) = 0.34, *p* = .73.

**Figure 2.**
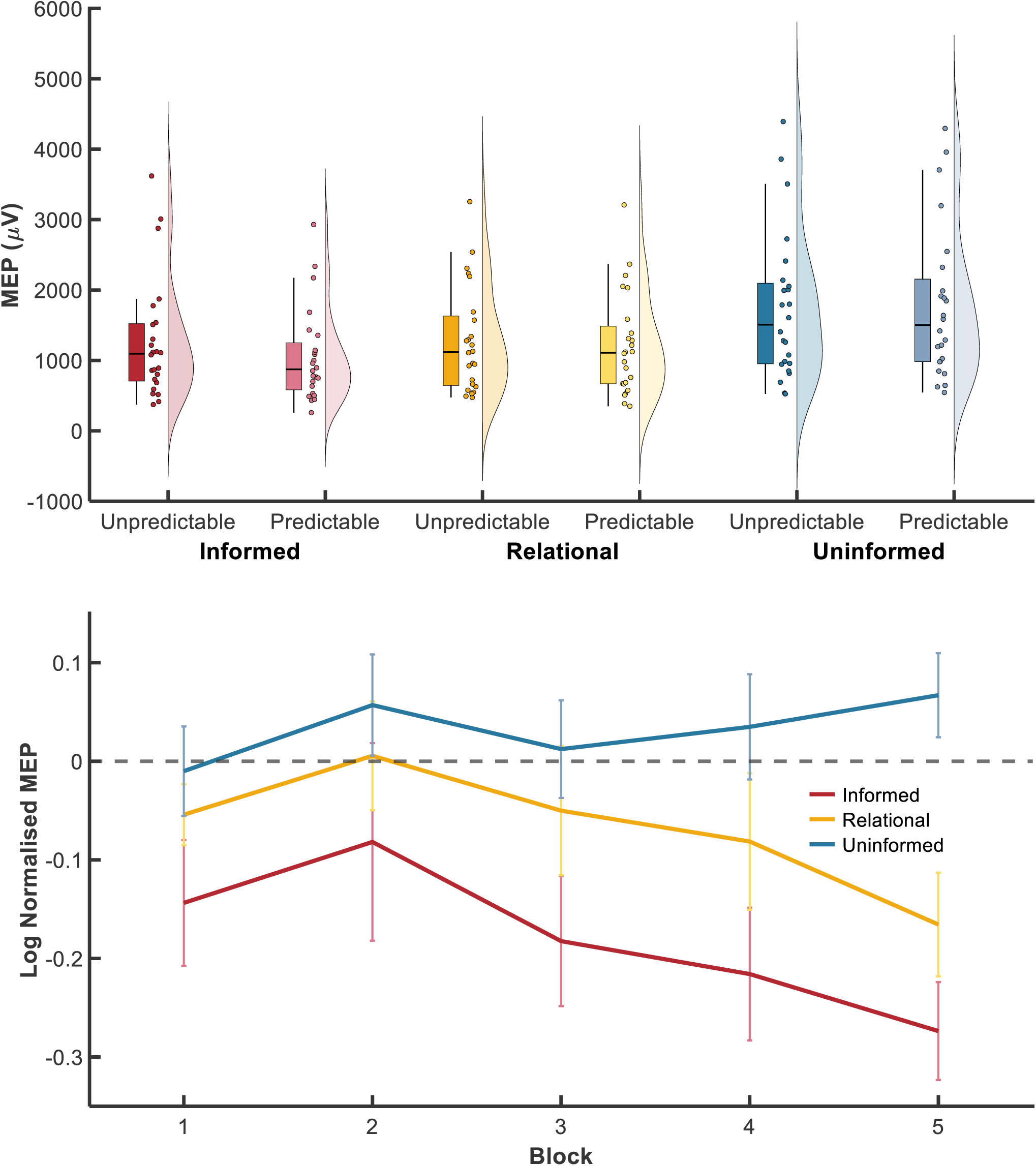
Raw MEPs collapsed over Block for unpredictable and predictable TMS by Instruction condition (top). Log normalised MEPs by Instruction condition and Block (bottom). MEP = motor-evoked potential.

**Table 1.** One sample t-test by Instruction condition and Block comparing log normalised MEPs against baseline.

|  |  | Block |  |  |  |  |
| --- | --- | --- | --- | --- | --- | --- |
|  |  | 1 | 2 | 3 | 4 | 5 |
| Informed | *t* | -2.25 | -0.82 | -2.77 | -3.21 | -5.51 |
|  | *p* | 0.03 | 0.42 | 0.01 | 0.004 | < 0.001 |
| Relational | *t* | -1.75 | 0.10 | -0.77 | -1.18 | -3.15 |
|  | *p* | 0.09 | 0.92 | 0.45 | 0.25 | 0.004 |
| Uninformed | *t* | -0.22 | 1.11 | 0.25 | 0.65 | 1.57 |
|  | *p* | 0.83 | 0.28 | 0.81 | 0.52 | 0.13 |

*Note.* Values < 0 indicate an attenuation effect with MEPs to TMS following a predictable cue less than MEPs to TMS following an unpredictable cue. MEP = motor-evoked potential; TMS = transcranial magnetic stimulation. We report the full data by condition and block: correcting for 3 comparisons (conditions) yields an alpha of 0.016, correcting for 5 comparisons (blocks) yields an alpha of 0.01, and correcting for 15 comparisons (condition × blocks) yields an alpha of 0.003.

Based on the post-experimental questionnaire, 51 participants were classified as Aware and 21 participants were classified as Unaware. Of the 21 Unaware participants, 2 were from the Informed condition, 6 were from the Relational condition, and the remaining 13 were from the uninformed group. Log normalised MEPs for Aware participants were below baseline for all blocks except block 2 (see Figure 3). For Unaware participants, log normalised MEPs remained around baseline for all blocks Table 2 includes the full list of one-sample t-tests by Awareness Classification and Block comparing log normalised MEPs against baseline (i.e., the attenuation effect). The contrast comparing contingency Aware versus Unaware showed that overall, across blocks, the Aware participants had significantly lower log normalised MEPs than the Unaware participants; *t*(70) = 3.112, *p* = .003. This contrast did not significantly interact with Block as a linear trend, indicating insufficient evidence that the effect of awareness on MEPs varied linearly across blocks; *t*(280) = 0.98, *p* = .33.

**Figure 3.**
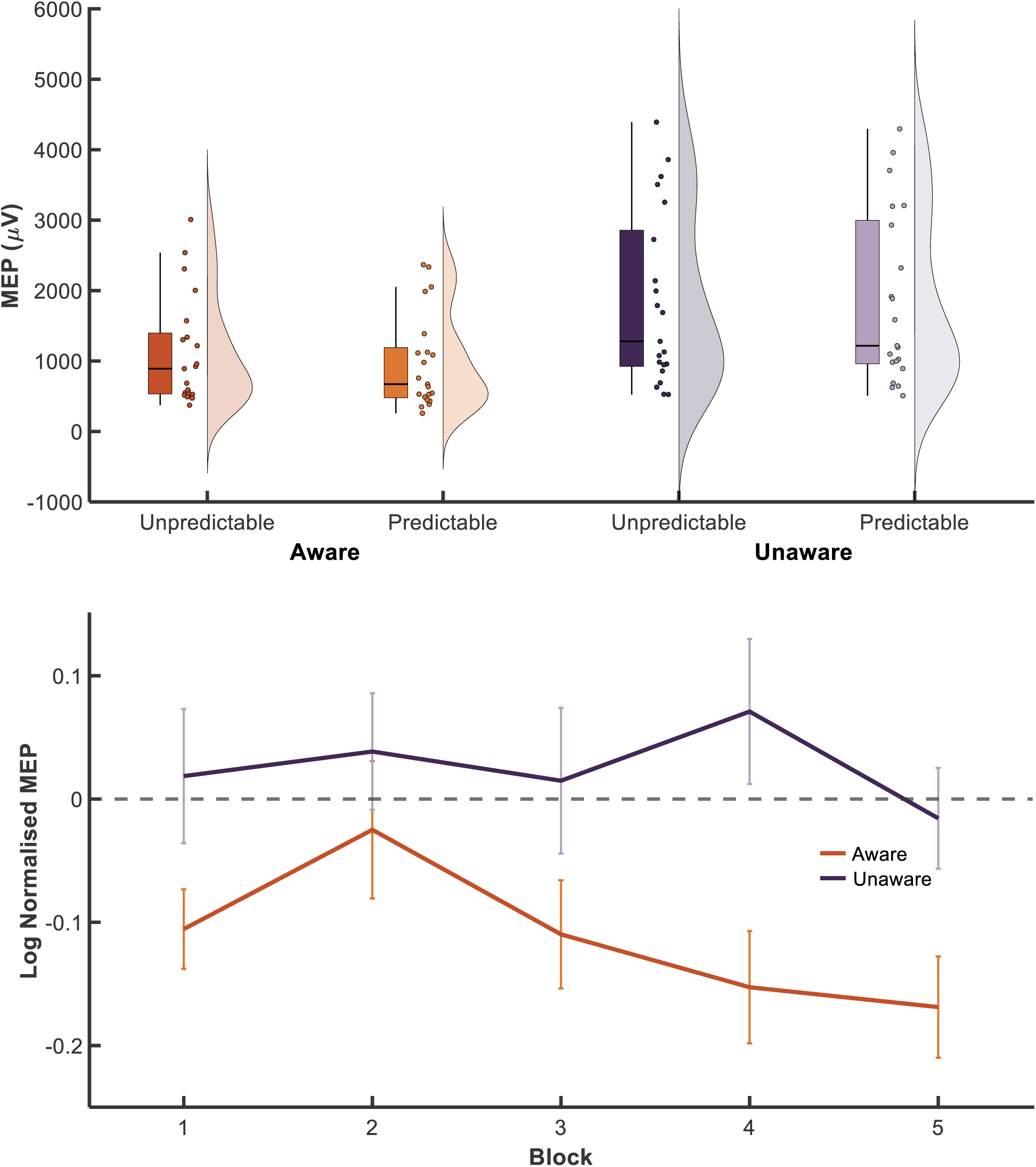
Raw MEPs collapsed over Block for unpredictable and predictable TMS by Awareness classification condition (top). Log normalised MEPs by Awareness classification and Block (bottom). MEP = motor-evoked potential.

**Table 2.** One sample t-test by Awareness classification and Block comparing log normalised MEPs against baseline.

|  |  | Block |  |  |  |  |
| --- | --- | --- | --- | --- | --- | --- |
|  |  | 1 | 2 | 3 | 4 | 5 |
| Aware | *t* | -3.26 | -0.45 | -2.50 | -3.36 | -4.11 |
|  | *p* | 0.002 | 0.66 | 0.02 | 0.002 | < 0.001 |
| Unaware | *t* | 0.34 | 0.82 | 0.25 | 1.21 | -0.38 |
|  | *p* | 0.74 | 0.42 | 0.80 | 0.24 | 0.71 |

*Note.* Values < 0 indicate an attenuation effect with MEPs to TMS following a predictable cue less than MEPs to TMS following an unpredictable cue. MEP = motor-evoked potential; TMS = transcranial magnetic stimulation.

## Discussion

Predictable TMS results in reduced motor system activity relative to unpredictable TMS. The current experiment tested whether this prediction-based motor attenuation can develop automatically, without conscious awareness of the predictive cue-stimulation relationship. We tested this hypothesis by masking the relationship between a warning cue and stimulation, presenting the warning cue embedded within a 1-back task, while delivering TMS embedded within a counting task. Participants who were informed and aware that TMS was reliably preceded by the warning cue and could anticipate the stimulation showed attenuated MEPs on predictable TMS trials relative to unpredictable trials. The central question of interest was whether participants who received reliable cueing of the warning cue-stimulation association, but were not informed or aware of this relationship, still showed differences in motor activity between predictable and unpredictable TMS.

First, we found that instructions about the predictive relationship affected the strength of motor attenuation. When participants were not informed of the relationship between the two tasks, masking the contingency between the warning cue and TMS was very effective. One possible mechanism for this masking is that participants compartmentalised the task into separate representations (see Duncan et al., 2020), making it difficult to notice or attend to the predictive relationship between the two tasks. Participants in this Uninformed group did not show a motor attenuation effect. The Relational group were instructed that there was a relationship between the two tasks but were not explicitly informed about the warning cue; these participants showed an increase in the strength of attenuation throughout the task, reflective of gradual learning of the association. When participants were explicitly informed about which cue reliably preceded TMS, they showed the greatest motor attenuation.

It is noteworthy that there was also an increase in the strength of attenuation for the Informed group across blocks. Although participants showed a significant attenuation effect from block 1, post-hoc analysis confirmed the there was a downward linear trend across blocks (*t*(92) = 2.25, *p* = .03). If explicit knowledge about the relationship was sufficient, in and of itself, to produce attenuation then one might expect attenuation to have reached its maximum from block 1. The results indicate that the attenuation effect strengthened over blocks, which can be interpreted in two ways. It is possible that this pattern of results in the Informed group reflect a combination of description and experience learning, whereby the attenuation in early blocks is driven by knowledge provided from the instructions, while the attenuation in the alter blocks are driven both by this knowledge and self-experienced learning. Alternatively, it is possible that participants had the required knowledge necessary for showing stronger attenuation in the early blocks but were unable to express this knowledge due to the complex nature of the dual task. The task requires continuous monitoring and responding of a visual n-back sequence while simultaneously counting TMS pulses embedded in a sequence with auditory tones. One way to test this second explanation would be to give participants practice with the dual task before introducing the critical warning cue-TMS relationship. In the current design, we gave participants practice with each task separately, before combining the two tasks and introducing the predictive relationship immediately. If the weaker attenuation effect in early blocks is due to the complexity of the dual task, practice on the dual task should reduce its interference on the attenuation effect.

Second, we assessed awareness by probing knowledge of the cueing relationship at the end of the experiment. Based on three questionnaire items, participants were classified as either Aware (mostly comprising Informed or Relational participants) or Unaware (mostly Uninformed participants). Participants who demonstrated contingency awareness of the predictive relationship showed strong motor attenuation for cued TMS, whereas those who remained unaware of this relationship by the end of the experiment did not show a motor attenuation effect.

Together, the instructional manipulation and the awareness classification results provide complimentary evidence that motor attenuation requires attention to the critical warning cues and knowledge of the cueing relationship. These findings suggest that prediction-based motor attenuation does not develop automatically and has important theoretical implications for predictive coding. There are mixed reports on whether prediction-based attenuation or expectation violations in the sensory system can develop automatically with little or no conscious awareness (e.g., Bekinschtein et al., 2009; Tivadar et al., 2021). Our current results support the view that, at least in the motor system, prediction-based attenuation and expectation violation cannot develop without awareness.

One possibility is that motor attenuation can occur automatically but that our paradigm was not amenable to detecting such effects. The dual task design requires attenuation to be allocated across the visual domain for processing the warning cue and the motor domain for processing the stimulation, and may be insufficiently sensitive for detecting automatic motor prediction errors. The cross-modal nature of the design, which involved a predictive relationship across visual and motor domains, may require more processing to integrate information compared to the processing of information within a single domain. Previous studies reporting automatic prediction effects in the sensory domain have used auditory oddball paradigms. Here, a repeating sequence of tones acts as a cue for an upcoming tone, and a tone that deviates from this predictable tone violates expectations. This oddball design requires information to be integrated within a single domain, where the stimulus to be predicted and the cues signaling the stimulus are within the same modality. Future studies may adopt similar methods in the motor domain to rule out the possibility that the failure to observe automatic prediction effects in the current experiment is due to information needing to be integrated across two domains. Reciprocally, future studies may adopt our current two task masking design in the sensory domain to test the role of non-conscious effects.

On the other hand, the current results may challenge the commonly held view that predictive coding mechanisms, specifically the updating of internal models via the propagation of prediction errors, is an automatic process. Most studies in support of the automaticity of prediction error signals have used classic oddball paradigms which do not control for adaptation to sensory signals within a sequence (e.g., Koelsch et al., 2006; Simpson et al., 2002), that is, the reduction in neural response to a sound, simply due to repeated presentation of that sound. Studies that have used a “local-global” design that controls for local stimulus predictability while manipulating a global expectation violation have found little evidence in support of automatic prediction effects in unconscious patients (e.g., Bekinschtein et al., 2009; Nourski et al., 2018). Consequently, the results of the current study, together with the evidence from studies using odd ball designs controlling for adaptation effects, add to the body of evidence indicating a necessary role of awareness for the detection of prediction errors.

In summary, the findings from the current experiment suggest a critical role for awareness in prediction-based motor attenuation, particularly when information needs to be integrated across domains. While previous studies using auditory oddball tasks have shown prediction effects in individuals with reduced consciousness, our study found no evidence of motor attenuation in participants who were unaware of the predictive relationship between a visual cue and a TMS pulse. This dissociation indicates that motor predictive processes may differ from the automatic principles governing sensory prediction. One possibility is that motor prediction relies more heavily on attention and top-down cognitive processing, or that the integration of information across domains requires conscious awareness, limiting the capacity for implicit learning. An alternative possibility is that awareness plays a more critical role in prediction-based attenuation, both in the motor and sensory system, than is typically claimed. Our results show a necessary role of contingency awareness for motor attenuation in a visual cueing paradigm and add to the growing evidence questioning the support for automatic prediction-based attenuation or unconscious detection of expectancy violation when confounding factors such as adaptation have been controlled.

## Author Contributions

D. T., and L. L. designed the research; L. L., and C. T. performed the research; D.T. supervised the research; D. T. analyzed the data; D. T., C. T., and E. L. wrote the paper

## Acknowledgements

The authors thank Monique Cost-Chretien and Hilary Don for their input on the manuscript.

